# Deep neural networks for interpreting RNA binding protein target preferences

**DOI:** 10.1101/518191

**Authors:** Mahsa Ghanbari, Uwe Ohler

## Abstract

Deep learning has become a powerful paradigm to analyze the binding sites of regulatory factors including RNA-binding proteins (RBPs), owing to its strength to learn complex features from possibly multiple sources of raw data. However, the interpretability of these models, which is crucial to improve our understanding of RBP binding preferences and functions, has not yet been investigated in significant detail. We have designed a multitask and multimodal deep neural network for characterizing in vivo RBP binding preferences. The model incorporates not only the sequence but also the region type of the binding sites as input, which helps the model to boost the prediction performance. To interpret the model, we quantified the contribution of the input features to the predictive score of each RBP. Learning across multiple RBPs at once, we are able to avoid experimental biases and to identify the RNA sequence motifs and transcript context patterns that are the most important for the predictions of each individual RBP. Our findings are consistent with known motifs and binding behaviors of RBPs and can provide new insights about the regulatory functions of RBPs.

## 1 Introduction

RNA-binding proteins (RBPs) play important roles in all aspects of post-transcriptional gene regulation including splicing, polyadenylation, transport, translation, and degradation of RNA transcripts [Gerstberger et al., 2014]. It is therefore not surprising that misregulation of RBPs as well as mutations in their protein sequence and/or their mRNA targets can result in diseases including cancer [Cooper et al., 2009, Siddiqui and Borden, 2012]. Hence, it is essential to identify RBP binding preferences to understand their function and reveal their disease promoting mechanisms. Although we are reaching a consensus annotation of all human RBPs [Ascano et al., 2012], and despite recent large-scale efforts [Van Nostrand et al., 2016], the binding preference and targets of comparatively few of these are well determined [Wheeler et al., 2018].

Crosslinking and immunoprecipitation followed by sequencing (CLIP-seq) protocols have made it possible to characterize transcriptome-wide binding sites of RBPs [Hafner et al., 2010, Konig et al., 2010, Van Nostrand et al., 2016]. Despite providing a valuable resource, CLIP data needs to be regarded with caution. Compared to alternatives such as RNA-binding and immunoprecipitation (RIP), CLIP results in significantly larger numbers of target sites, indicating possible cross-linking of low-specificity events, or that only few mRNA copies of a given gene are actually bound in the same cell [Mukherjee et al., 2011, Plass et al., 2017]. On the other hand, CLIP-seq is sensitive to expression levels, meaning that binding events on lowly expressed transcripts may not be detected. Finally, CLIP protocols are variable, and aspects of the protocol can introduce significant biases, most notably by the type and concentration of RNase that is used [Kishore et al., 2011]. To derive binding sites from CLIP-seq reads, several specialized peak detection methods have been developed to capture high-fidelity RBP binding sites from different CLIP protocols [Corcoran et al., 2011].

Motif finding approaches can extract the dominant shared sequence/structure motifs that characterize the binding sites, ranging from those based on sequence only [Bailey, 2011, Georgiev et al., 2010] to more recent ones that also take aspects of RNA structure into account [Kazan et al., 2010, Heller et al., 2017, Munteanu et al., 2018]. These approaches aim at deriving optimal continuous sequence/structure motifs based on an e.g. information theoretic objective function. Alternatively, binding sites can also be analyzed by classification approaches, for instance to distinguish between bound and unbound sites. Models with this aim use large numbers of binding sites (and possibly their flanking regions), typically for one RBP in one cell type at a time. The trained model can then be used to reveal missing targets of the RBP in the specific cell type, or to identify putative target sites that are bound in other cell types without in vivo binding data [Maticzka et al., 2014, Stražar et al., 2016]. However, interpreting these classifiers, e.g. to derive consensus motifs as in motif finding, is usually not straightforward.

The rise of deep learning has spurred the recent development of deep neural network (DNN) to predict TF or RBP binding sites. Alipanahi et al. first showed that convolutional neural networks (CNNs) can learn TF/RBP binding sites with high accuracy compared to state of the art methods by using only the DNA/RNA sequences as input. Since then, several convolutional and recurrent neural network models for genomics data have improved prediction accuracy [Quang and Xie, 2016]. For example, iDeep [Pan and Shen, 2017] leverages a multimodal DNN to integrate different sources of data to infer RBP binding sites. A study concurrent to ours additionally included relative distances of binding sites to various positional landmarks such as splice sites, using spline transformations [Avsec et al., 2018].

While these deep models show great promise to push the accuracy of predictions, it is generally unclear what the models base these predictions on. Using CNNs with sequence as input makes it possible to inspect the kernels or convolutional filters in the first DNN layers. One can extract the weights of these kernels or aggregate input subsequences that maximally activate the kernels and visualize them as position weight matrices (PWMs) [Pan and Shen, 2017, Alipanahi et al., 2015, Avsec et al., 2018]. While these patterns give general insight about low-level representations that the model has learned, they do not provide information about the decision itself, especially for DNNs with multiple layers. This challenging problem of explaining predictions has become an active field of study, and several methods have been developed over the last couple of years [Shrikumar et al., 2017, Lanchantin et al., 2016, Sundararajan et al., 2017].

In this work, we propose DeepRiPe (Deep RBP binding Preference), a multitask and multimodal DNN model set up with the aim to characterize RBP binding preferences. DeepRiPe uses a modular structure to learn informative features from DNA sequence and transcript region types, as many RBPs have preferences for binding to specific regions of a transcript. We frame RBP site prediction as multitask learning problem, i.e. predicting binding sites for several RBPs simultaneously. This allows the model to use shared information among tasks; as we well show, this is critical to focus the model on the distinctive features of RBP binding sites: it avoids putting strong weights on generic features of binding sites across tasks (e.g. UTR sequence composition, accessible secondary structure) as well as protocol inherent biases, e.g. resulting from sequence preferences of RNase enzymes. In turn, several RBPs may possess similar binding patterns, and sharing information among their predictor can help the model when training data is limited. We evaluate DeepRiPe on a large compendium of PAR-CLIP datasets and use Integrated Gradients (IG) to interpret the model [Sundararajan et al., 2017]. Investigating different model choices, we show that a seemingly good performance does not necessarily imply that the model has learned biologically relevant features.

## 2 Methods

### 2.1 Model design and training

In this work, we use two types of DNN architectures, Convolutional Neural Networks (CNNs) and Recurrent Neural Networks (RNNs) [Goodfellow et al., 2016]. CNNs use a weight-sharing strategy and they are highly successful to locate motifs, for example in a sequence, independent of their position within the sequence. RNNs have a “memory” that allows information to persist so that they can learn dependencies in the data. More specific, we use a bidirectional gated recurrent network (GRU) [Chung et al., 2014] to account for possible long-range dependencies of the features.

The model consists of a sequence module that extracts features from the RNA sequence and a region module that extracts features from genomic locations. The features of these modules are then merged and fed to a multitask module to predict the binding sites of multiple RBPs simultaneously. Figure 1 shows a simplified architecture of the model.

**Figure 1:**
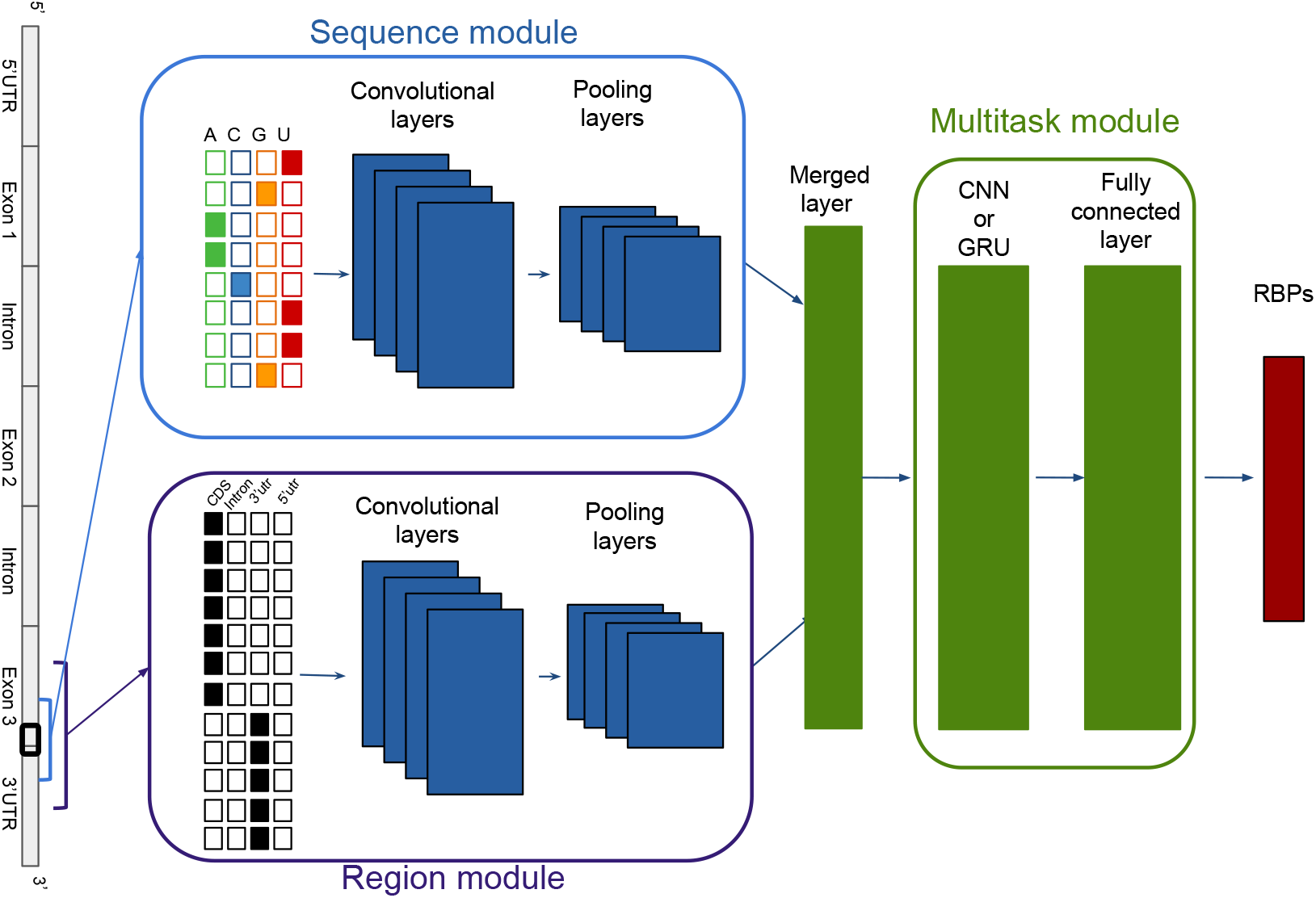
A simplified graphical illustration of the model: the model consists of a sequence module that extracts features from the RNA sequence and a region module that extracts features from genomic locations. The features of these modules are then merged and fed to a multitask module to predict the binding sites of multiple RBPs simultaneously.

The sequence and region modules both have a convolution layer followed by a rectified linear unit (Relu), a max pool layer, and a drop out layer with probability of 0.25. We used 90 filters with length 7 for both convolution layers. The multitask module takes as input the concatenated features from sequence and region modules and consists of one CNN (with 100 filters of length 5) or one bidirectional GRU (with 60 units) and one fully connected layer with 250 hidden units and Relu activations. The output layer contains k sigmoid neurons to predict the probability of binding, one for each RBP.

To assess the contribution of different aspects to the success of the DNNs, we also explore variations of the architecture and training of the model; note that in singletask models, where the model predicts the binding sites of one RBP, the output layer has only one neuron. In all applications, we use CNNs in the multitask module unless stated otherwise. The training was performed with an Adam optimizer [Kingma and Ba, 2014] using a minibatch size of 128 for 20 epochs to minimize the mean multi-task binary cross entropy loss function on the training set. To account for imbalanced data, we used a weighted loss function which gives higher penalties for misclassifying samples related to the classes with less samples. The best model was chosen based on validation loss computed at the end of each epoch. We used early stopping to prevent the possibility of over-fitting during the training.

### 2.2 Input data

We collected PAR-CLIP datasets for 59 RBPs from different publications, which were profiled with the same flag-tagged construct in HEK293 cell line. These libraries were quality controlled and processed with the same pipeline, including PARalyzer [Corcoran et al., 2011] for peak calling and the human GRCh37/hg19 release as reference, in a recent study [Mukherjee et al., 2018]. We chose RBPs that have between 1000 and 10^6^ peaks and divided them into 3 categories: RBPs with more than 10^5^ peaks, RBPs that have between 15000 and 10^5^ peaks and RBPs with less than 15000 peaks. We used RBPs in each category for training and evaluating a separate DNN which we refer to model-high, model-mid and model-low respectively. More details about the data can be found in Supplementary Material(SM) Section 1.

To prepare the data for input to the DNN, we first split the genome into 50-bp non-overlapping bins and kept only bins that overlap with the transcriptome. For each bin, we assign a label vector with k entries correspond to all RBPs of interest to define the labeled data for the multitask model. For each bin, the label of an RBP is 1 if more than half of its peak region falls within a 50-bp bin and 0 otherwise. We kept only bins with at least one binding event. In this way, the negative samples of one RBP may serve as positive samples of other RBPs. We used 20 and 10 percent of the bins for validation and test of the model respectively and the rest of the bins for training the model. Note that all of the results presented in this study are obtained using independent test data.

The input for the sequence module is one-hot encoded DNA sequence from a 150-bp sequence; the input for the region module is one-hot encoded region features from a 250-bp sequence, both centered on each 50-bp bin. While sequence features denote the nucleotide (A,C,G,U), the region features annotate each position as being in a 3’UTR, 5’UTR, CDS, or intron region. The flanking regions can give insight about the context of binding sites, and by using single nucleotide resolution, we can capture whether binding sites occur at boundaries of region types (e.g. exon/intron junctions, cleavage sites) near crosslinked sites.

To evaluate the model on data obtained from other protocols and cell lines, we also applied it on eCLIP [Van Nostrand et al., 2016] processed binding sites (intersection between two replicates;) provided by ENCODE website (https://www.encodeproject.org) and predict binding for them using DeepRiPe trained on PAR-CLIP. To define comparable input vectors for DeepRiPe, we extended the middle of each eCLIP peak with 75 bp and 125 bp both upstream and downstream for sequence and region modules respectively.

### 2.3 Evaluation scores

We evaluate the DeepRiPe model, which is trained using training and validation sets, on independent test data. Classification performance is assessed by both the receiver operating characteristic (ROC) and precision-recall (PR) curves, as well as the area under the ROC curves (denoted as AUROC). Average precision (AP) summarizes PR curves and is defined as the precision averaged across all values of recall. AP is more conservative compared to area under the PR curve since the latter uses linear interpolation and can be too optimistic. Note that AP is more appropriate than AUROC in the case of imbalanced data with more negative samples, since it does not take into account the number of true negatives.

### 2.4 Interpretation

While obtaining accurate predictions of RBP/TF binding sites is important, it is at least equally important, for instance for an eventual application to interpret non-coding genetic variation, to understand why the model makes these predictions and which parts of the input contribute the most to the output. The gradient (partial derivatives) of an output neuron with respect to its input indicates how the output value changes with respect to a small change in inputs. This is the basic concept used in gradient-based attribution methods that assign an attribution value to each input feature of the network, indicating how much that feature contributes to the output. Here, the target neuron of interest is the output neuron associated with the corresponding RBP class for a given sample, and an attribution method can specify which nucleotides of the sample input sequence and/or which region part were responsible for the output of the RBP. In this study, we use an attribution method called Integrated Gradients (IG) [Sundararajan et al., 2017]. IG computes the average gradients of the output as the input varies along a linear path from a baseline or reference to the input, to avoid the saturation problem that occurs when computing gradients only at the input. The baseline is defined based on the application and often chosen to be zero. We used zero and 0.25 for the baselines of sequence and region inputs respectively.

Calculating all the attribution values corresponding to all positions of one input sample leads to an “attribution map” of the sample. By visualizing the attribution map as sequence logos (for sequence) or barplots (for region), we can observe the influence of each position (for example a nucleotide of a specific input sequence) on the prediction. The height of sequence logos or barplots indicates the importance of that position in the prediction. Positions with large positive attribution values can be interpreted as features that were informative for the prediction of the RBP. Visualization of the attribution maps of each input sample for a specific RBP not only reveals the potential target motif or motifs of the RBP, but it can also be used to locate the potential binding sites of the RBP on a new sequence or to assess the effect of genetic variants on RBP binding site.

To obtain consensus motifs for each RBP we aggregate the results of all attribution maps corresponding to all binding sites with high prediction scores (prediction score > 0.5). We reason that high confidence binding sites most probably contain the target motifs, while the ones with low probability may result from spurious binding. The detailed description of the method can be found in SM Section 4.

## 3 Results

### 3.1 Results of the model

#### 3.1.1 Performance

The main goal of our study is to establish interpretable classifiers, as a first step towards models that can quantify the impact of sequence variation on post-transcriptional gene regulation. To start, we established the baseline performance of DeepRiPe using ROC and precision-recall(PR) curves. To account for the drastic differences in the number of called peaks (ranging from approximately 1,000 – 1,000,000 sites), DeepRiPe consists of three networks with identical architectures (Figure 1), each of which is trained on a subset of CLIP datasets with comparable binding site numbers (see Methods). Figure 2A shows the the ROC and PR curves as well as the corresponding AUROC and AP values for a subset of 15 RBPs, which we investigate in more detail below. The AUROC and AP values for all RBPs are provided in SM Section 2. While all AUROC values are above 0.7, the AP scores exhibit a wide range. Figure 2B shows the distribution of prediction scores for 3 RBPs, where two of them (MBNL1 and ORF1) have high AUROC and AP scores and the third (CPSF6) lower scores. Consequently, while there is a clear difference between the distributions of the scores for positive and negative samples for MBNL1 and ORF1, this is less clear for CPSF6.

**Figure 2:**
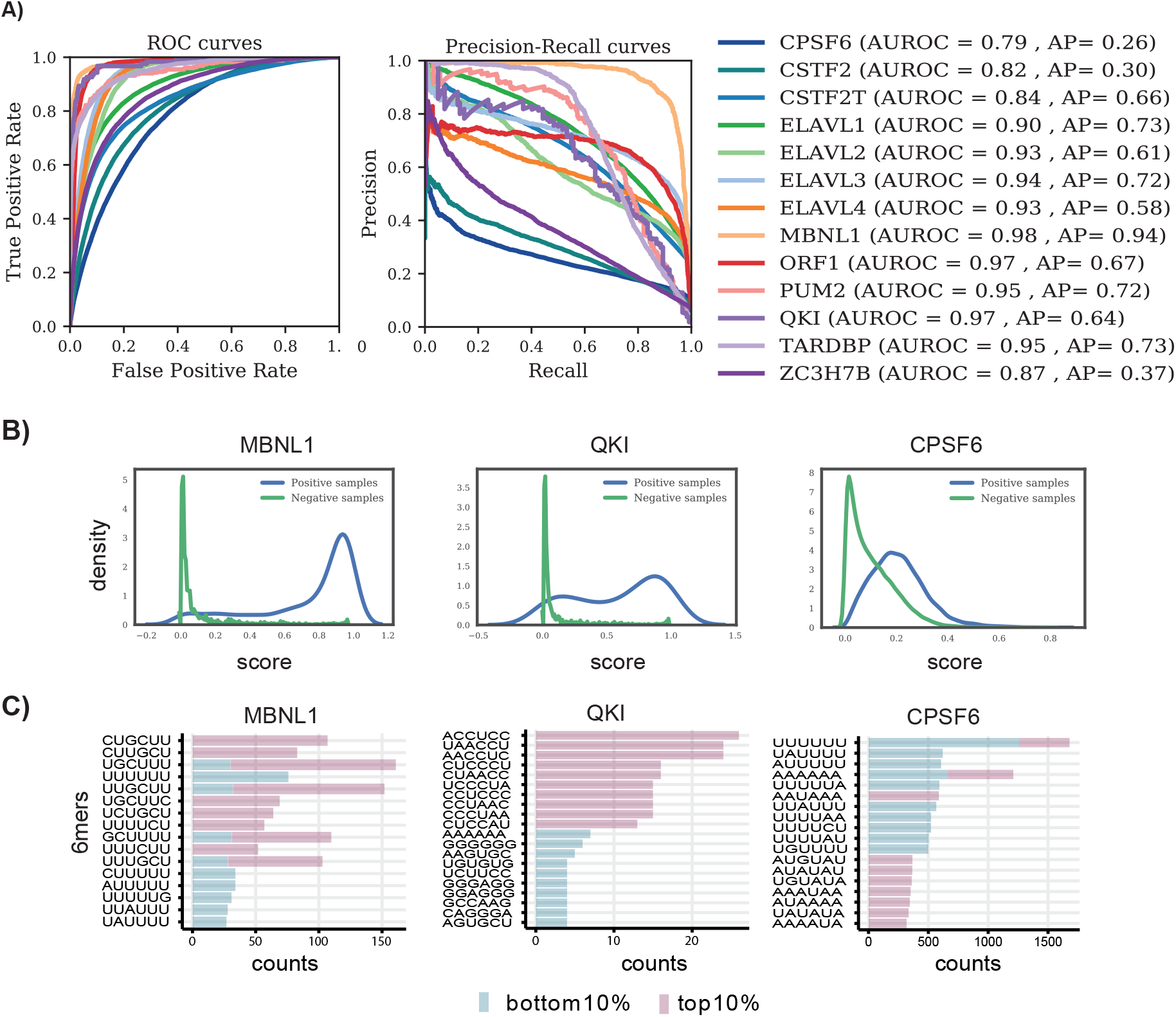
A) ROC and Precision recall curves for several RBPs. The corresponding AUPRC and AP scores are shown in parentheses. B) Prediction scores distribution for positive and negative samples for several RBPs and 6mers counts at the top and bottom 10 percent of positive samples ranked based on their prediction scores.

Variation in network performance can result from the different quality of individual CLIP datasets, which may on the one hand miss genuine binding sites (false negatives), and on the other hand contain substantial amounts of false positive, low-affinity cross-linked sites. To investigate this, we ranked the CLIP peaks, i.e. the candidate binding sites of each RBP (positive samples of test data), based on the prediction score and extracted the 6mers from the bottom 10 percent as well as the top 10 percent of the sites. Figure 2C shows the top 10 6mers in each set for four RBPs. While the top 6mers in the high ranking sites are in line with the corresponding RBP motif(s), this is not necessarily the case for low ranking sites specially for RBPs with low scores. As an example for CPSF6 with AP scores of 0.26 the high ranking binding sites contains mostly AAUAAA and UGUA elements while the low ranking sites are enriched in U-rich elements that have been previously reported as CLIP artifacts [Krakau et al., 2017]. This suggests that the RBP data used in our study may vary considerably in their quality, and indicates a potentially high rate of false positives in some of the (PAR-)CLIP datasets. The aim of our study is therefore not to achieve the best performance according to some metric; simply striving for classification performance can be highly misleading if the data is subject to considerable biases.

#### 3.1.2 Interpretation

The results so far emphasize the need for an interpretable classifier, to better understand what the driving input features are behind a good or poor performance. To this end, we applied methods that provide model interpretability, to determine which sequence and region type patterns are informative for predicting RBP binding sites. For each RBP, we computed attribution maps for positive samples of test data with the highest prediction scores from the model. For any given input sequence, an attribution map highlights the individual nucleotides, respectively the relative position and region type, that were most important for its classification as target site for a specific RBP. Figure 3 shows attribution maps for target sequences of several RBPs. The examples illustrate that the model is able to learn and highlight important sequence motifs present in the input; these motifs in fact agree with the known motifs for that particular RBP.

**Figure 3:**
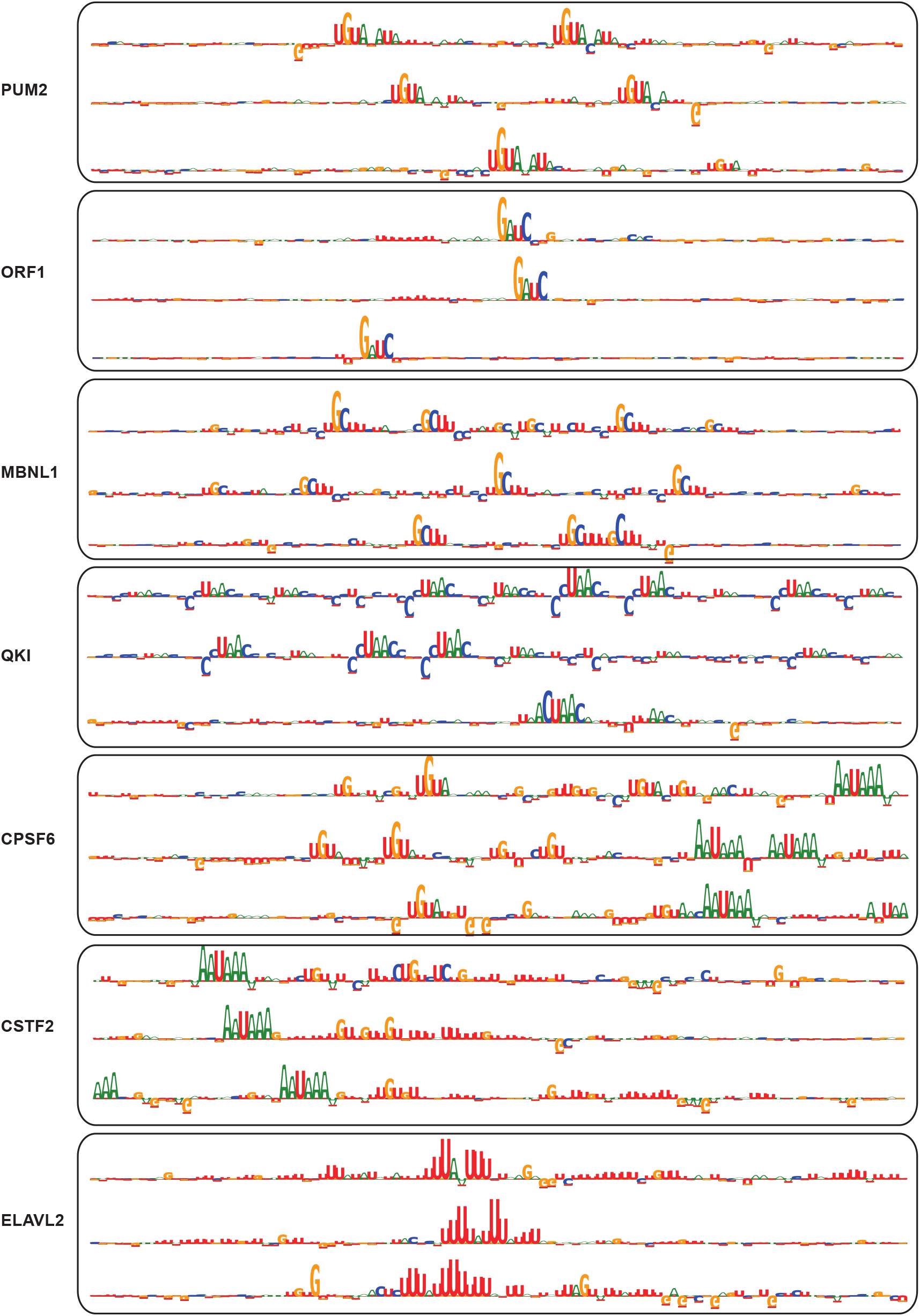
Interpretation of the model using attribution maps obtained from IG method: The sequence logos corresponding to attribution maps for true binding sites of corresponding RBPs with the highest DeepRiPe prediction scores.

Looking at specific factors in more detail highlights a crucial advantage of DNNs for regulatory sequence interpretation: the models are able to locate both simple and complex patterns in the input, i.e. one to several occurrences of a motif and also composite motifs, without additional prior knowledge. As examples for simple patterns, we observe the well-established UGUAHAUA binding motif in attribution maps corresponding to PUM2. ORF1 is a protein encoded by the transcripts of LINE-1 retrotransposable elements and responsible for its retortransposition; attribution maps of its target sequences delineate with high precision its GAUC target motif [Mandal et al., 2013].

MBNL1 and QKI are splicing factors with reported YGCU/GCUU [Lambert et al., 2014] and ACUAAY [Hafner et al., 2010] binding motifs, respectively, and their attribution maps reveal several occurrences of the motifs in the mRNAs. ELAVL2, ELAVL3, ELAVL4 are RBPs that regulate mRNA stability and translation through the 3’UTR and bind to U-rich elements [Keene, 2001]. The patterns observed in their attribution maps are consistent with this knowledge. ELAVL1 additionally binds to pre-mRNAs in the nucleus and thus to additional region types, and also showed similar preference for U- and AU-rich patterns [Keene, 2001]. Depending on the input sequence, we are also able to identify variable numbers of the core U-rich pentamer.

The model is also able to locate combinations of motifs. For example, we observe RNA polyadenylation/cleavage related sequence elements, namely AAUAAA and U/GU-rich elements, located upstream and downstream of the actual site of cleavage [Darmon and Lutz, 2012] respectively, in the attribution maps of cleavage and polyadenylation specificity factors (CPSFs) and cleavage stimulatory factors (CSTFs), respectively. Moreover, the previously reported motif UGUA is observed in attribution maps of CPSF6, which is involved in 3-end cleavage of RNA transcripts [Yang et al., 2011, Brown and Gilmartin, 2003].

To obtain consensus representations for each RBP, we aggregated the attribution maps corresponding to all binding sites with prediction scores larger than 0.5 as described in Section 2.4. The results are shown in SM Section 4. Altogether, patterns obtained from attribution maps were remarkably consistent with previously reported motifs, in spite of not optimizing an objective function that directly quantifies the presence of common, strong motifs as in traditional motif finding. It also adds confidence that the model has learned genuine sequence features. Notably, the DNN enables us to see the actual occurrence of the motif in the sequence, and it is also intrinsically able to identify complex motif patterns such as combinations of motifs, which is observed in the attribution maps of CPSF6 and CSTF2. This flexibility is also apparent in the frequent multimeric, closely spaced or adjacent occurrences of the consensus ELAV pentamer. This characteristic inherent flexibility of the DNN is a clear advantage over classical regulatory sequence analysis, with its rich literature of highly specific approaches for complex motif configurations.

#### 3.2 Benefits of the multimodal model

To assess the benefit of the multimodal model that uses both sequence and region type as input, and to evaluate the impact of region type in the performance of the method, we trained the DeepRiPe model without using region type information as input. Both sets of models were trained with similar structure and the same hyperparameters. Figure 4 shows the scatter plot comparing the AUROC and AP scores of the two models, as well as attribution maps corresponding to region input for ELAVL2 and CSTF2. The multimodal model using both sequence and region type outperforms the model that uses only sequence for nearly all of the RBPs. This indicates the importance of region type for prediction. The attribution maps also revealed that the model uses specific region preferences that are consistent with current knowledge. For example, functional ELAVL2 binding sites are predominantly located in the 3’UTR regions, and CSTF2 binds to the 3’ end of the gene.

**Figure 4:**
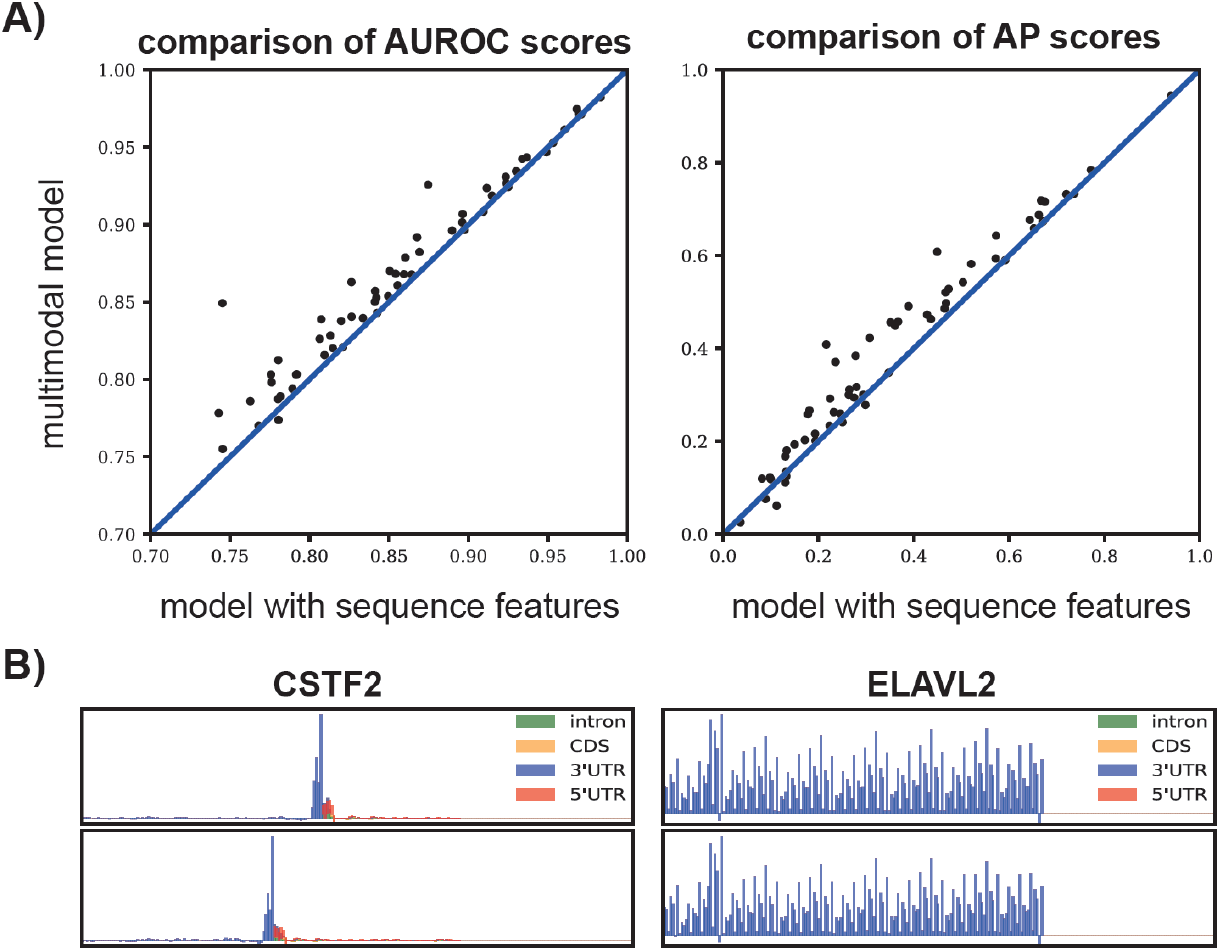
Assessing the performance of multimodal model. A) Scatter plots comparing the AUROC and AP scores of DeepRiPe and singlemodal (the model using only sequence features). Each data point represents an RBP and it falls above the diagonal when DeepRiPe outperforms the singlemodal model, B) Attribution maps correspond to region inputs obtained from multimodal and singlemodal using IG method for positives samples of ELAVL2 and CSTF2

#### 3.3 Benefits of multitask learning

To assess the benefit of multitask learning, we compared the results of the model to those obtained by its singletask counterparts, for which we used the same hyperparameters as for the multitask model. We evaluated multiple strategies to define singletask training data: In the first strategy (single models 1), we oversampled from positive samples of the training and validation datasets for each RBP to ensure an equal number of positive samples as negative samples in these datasets. In the second strategy (single models 2), we used random negative samples obtained from unbound transcripts for each RBP. We compared the performance and interpretability of two approaches and present the results in Figure 5. Finally, we also subsampled from negative samples of the training and validation datasets to ensure an equal number of negative samples as positive samples in these datasets (single models 3); results can be found in SM Section 5.

**Figure 5:**
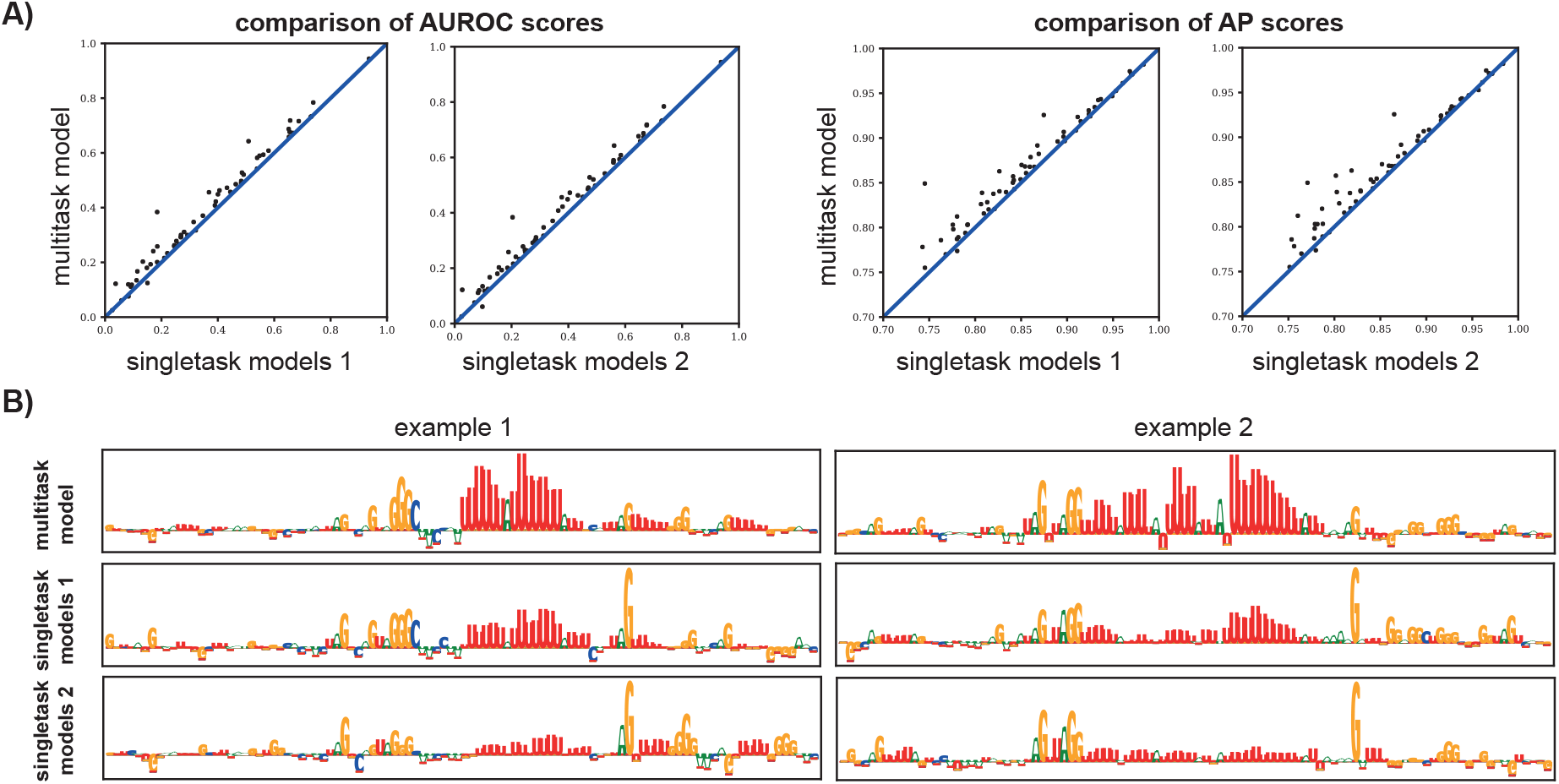
Assessing the performance of multitask model, A) Scatter plots comparing the AUROC and AP scores of DeepRiPe and singletask models. Single models 1 and single models 2 are trained on random negative samples from binding sites of other RBPs and unbound transcript respectively. Each data point represents an RBP and it falls above the diagonal when DeepRiPe outperforms its singletask counterpart, B) Comparing attribution maps obtained from multitask and singletask models using IG method for positives samples of ELAVL2.

The overall results indicate that for some RBPs, the multitask learning indeed boosts the performance. In a performance comparison to each of the three DeepRiPe submodels (model-high, model-mid and model-low; SM Section 5), it became apparent that RBPs with low number of samples benefit the most, which is in line with the promise of multitask learning.

While there is consistent but limited performance improvement between single-and multitask models, the interpretability of single- and multitask models differed considerably. Comparison of attribution maps of ELAVL2 (Figure 5B) revealed that the singletask models showed reduced importance of the known motif and were heavily misled by the PAR-CLIP sequence bias from RNase T1, which cleaves after guanines and is very prominent in especially early PAR-CLIP data sets [Kishore et al., 2011]. While the strategy of using binding sites of other RBPs as negative samples (single models 1) rather than using random negatives from unbound transcript (single models 2) leads to a slightly better delineation of the target motif, the multitask learning approach can reveal the actual motif clearly: When learning the preferences of multiple RBPs simultaneously, the cleavage bias does not constitute useful information to discriminate between target sites of different RBPs, as many PAR-CLIP peaks will be equally affected by it. Multitask learning thus puts much less weight on protocol biases that are shared between several RBP libraries.

#### 3.4 Generalization power of DeepRiPe

DeepRiPe as a classification method should be able to distinguish between bound and unbound sites for a specific RBP regardless of experimental conditions and therefore to identify putative binding sites in other cell types for which there is no PAR-CLIP data. To assert this ability to generalize, we used 6 eCLIP datasets of RBPs, which were profiled by both eCLIP and PAR-CLIP in different cell lines, namely CPSF6, CSTF2T, CSTF2, PUM2 and QKI (two additional cell lines) [Van Nostrand et al., 2016]. We ran the PAR-CLIP trained models on eCLIP targets, ranked eCLIP peaks for each RBP based on their DeepRiPe prediction score and counted all possible 6mers in the top 2000 (high confidence) and bottom 2000 (low confidence) binding sites. Figure 6 shows the top 10 6mers in each set. While top 6mers in high confidence binding sites resemble the motif(s) for the specific RBP, this is not the case for low confidence binding sites. As we observed on PAR-CLIP data, low-scoring eCLIP peaks are therefore likely to represent weak affinity or spurious binding sites.

**Figure 6:**
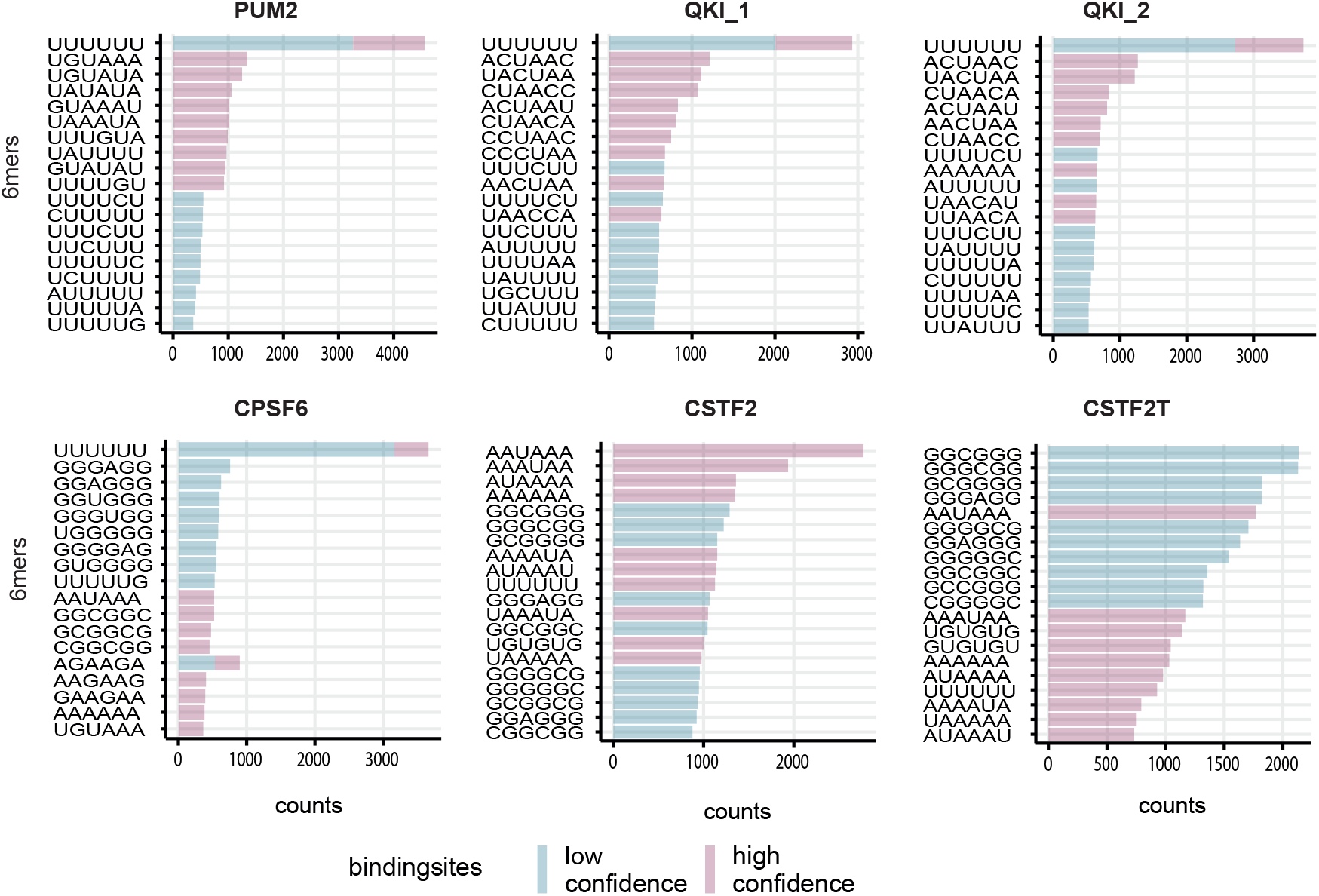
Performance of DeepRiPe on eCLIP data conducted in different cell types: top 6mers in high confidence as well as low confidence binding sites based on the prediction scores obtained from DeepRiPe.

#### 3.5 DeepRiPe as a potential tool to study the effects of sequence variants

Finally, we explored the potential use of DeepRiPe as a tool to identify and score sequence variants with potential impact on RBP binding. To this end, we used the trained model to compute and compare the attribution maps of wild-type and mutated sites for two RBPs.

ELAVL1 binds to the 3’UTR of the ERBB-2 oncogene mRNA. In a recent study [Epis et al., 2011], ELAVL1 was shown to oppose the repression effect of microRNA miR-331-3p in ERBB-2 by binding to a U-rich element (URE) near the miRNA target region. Mutation of the URE results in an experimentally detected shift of ELAVL1 binding to an upstream site with reduced binding affinity and weakens the repressive effect of ELAVL1 on miR-331-3p. Figure 7 shows the wild-type and mutant sequences as well as the resulting attribution maps. The attribution maps demonstrate the loss of ELAVL1 binding site at the mutated site while upstream sites were not affected, in line with the reported observation.

**Figure 7:**
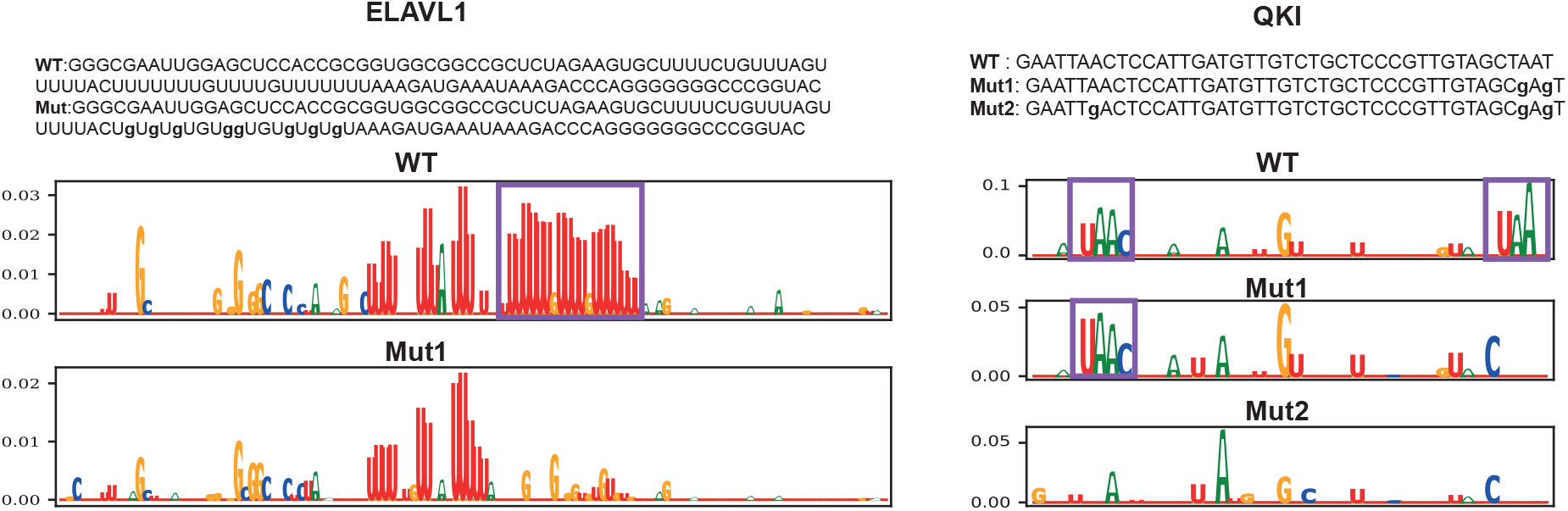
Sequences of wild-type and mutant constructs where mutations are shown in bold lowercase letters and their corresponding attribution maps for ELAVL1(A) and QKI(B). Abbreviation: WT, wild-type; Mut, mutant.

As a second example, we examined the effect of mutations in potential QKI binding sites in NUMB pre-mRNA. In a study that investigated the role of QKI in regulating NUMB alternative splicing [Zong et al., 2014], two mutant sequences Mut1 and Mut2 were generated targeting two potential binding sites of QKI in the regions surrounding the 3’ splice site of intron 12. While Mut1 contains mutations only in the second binding sites, Mut2 has mutations in both sites (Figure 7). Compared to wildtype RNA, with binding affinity comparable to that of a control RNA that carries a bipartite QKI consensus sequence, Mut1 RNA exhibited reduced QKI binding, while Mut2 RNA lost QKI binding completely. Figure 7 shows the attribution maps of the wild-type as well as the two mutated sequences. The attribution map of the wild-type sequence reveals a strong binding for the second binding site and a weak binding for the first binding site, the attribution map of Mut1 has lost the strong binding but preserves the weak binding, and the attribution map of Mut2 has lost both binding sites. These observations are consistent with the experimental findings.

## 4 Conclusion

We have developed a multimodal and multitask deep learning approach to model genuine, specific RBP binding events, and to extract informative features about RBP binding characteristics from dozens of high-throughput, noisy CLIP-seq data. The model recovers known sequence motifs and provides insight about RBP binding preferences. It can also locate the sequence motifs along the input sample and identify co-occurrences of motifs in a flexible manner. We observed considerable variability of success across different RBPs, and we were able to relate this to the absence of known motifs in low-scoring peaks; CLIP-seq experiments can result in tens of thousands of peaks, and it is highly unlikely that all of these represent targets with defined functional consequences of binding. Rather, large numbers of peaks may reflect poor antibody quality; sequencing artifacts; or interaction patterns of RBPs beyond specific sequence/structure target site definitions, such as helicases. As many peak callers do, our model assumes site-specific binding, and for libraries for which this assumption holds true, we are moving closer to a scenario in which we can now use the model to judge the quality of experiments, rather than to take noisy data as “ground truth”.

Singletask and multitask models solve different classification problems. While the overall performance of multitask and singletask reported here appear superficially similar, the multitask formulation of learning allows the model to focus on the features that are shared across the tasks. In this way, it is able to ignore possible protocol-inherent biases, as these will be present in datasets across different RBPs. We illustrate that this leads to notable differences in the features that a model utilizes for its predictions, with the multitask models relying more strongly on the presence of known motifs compared to the singletask methods. Choosing negative samples for each RBP from binding sites of other RBPs makes the prediction task harder, but at the same time it guides the model to learn specific motifs. Most previous RBP target classification approaches have been set up as singletask problems, which means that we cannot directly benchmark against them. In turn, many singletask models have been evaluated on cross-validated, held-out data from the same experiment. For some of these, the reported results will likely beoverly optimistic — the models will not generalize well, as we have recently observed anecdotally [Munteanu et al., 2018].

We have presented a deep network to extract insights about RBPs but can be extended in further studies. The method provides a tool to locate binding sites, and we anticipate that our strategy of using attribution maps to pinpoint motif occurrences can be extended to fully fledged motif finding. This holds promise to alleviating some shortcomings of current approaches for RBP motif discovery that struggle because of the shortness of the binding motif and the potentially large number of false positives in the input data. In this context, interpreting DNNs may provide competitive flexibility, as there is no need to specify parameter like motif length or configuration. For both classification and prediction, future work should address how to adequately consider RNA structure within the framework of deep neural networks, to advance the interpretation of non-coding sequence variants.

## Data and Software Availability

The code for DeepRiPe is available at https://github.com/ohlerlab/DeepRiPe.

## Acknowledgements

The authors would like to acknowledge Neelanjan Mukherjee for providing the processed PAR-CLIP data.

## Authors’ contributions

M.G. and U.O. conceived the project; M.G. developed the methodology with contributions by U.O. and implemented the method and performed the analysis. M.G and U.O. wrote the paper.

